# Stress relaxation amplitude of hydrogels determines migration, proliferation, and morphology of cells in 3-D

**DOI:** 10.1101/2021.07.08.451608

**Authors:** Jonas Hazur, Nadine Endrizzi, Dirk W. Schubert, Aldo R. Boccaccini, Ben Fabry

## Abstract

The viscoelastic behavior of hydrogel matrices sensitively influences the cell behavior in 3D culture and biofabricated tissue model systems. Previous reports have demonstrated that cells tend to adhere, spread, migrate and proliferate better in hydrogels with pronounced stress relaxation. However, it is currently unknown if cells respond more sensitively to the amplitude of stress relaxation, or to the relaxation time constant. To test this, we compare the behavior of fibroblasts cultured for up to 10 days in alginate and oxidized alginate hydrogels with similar Young’s moduli but diverging stress relaxation behavior. We find that fibroblasts elongate, migrate and proliferate better in hydrogels that display a higher stress relaxation amplitude. By contrast, the cells’ response to the relaxation time constant was less pronounced and less consistent. Together, these data suggest that it is foremost the stress relaxation amplitude of the matrix that determines the ability of cells to locally penetrate and remodel the matrix, which subsequently leads to better spreading, faster migration, and higher cell proliferation. We conclude that the stress relaxation amplitude is a central design parameter for optimizing cell behavior in 3-D hydrogels.

## Introduction

Culturing cells in 3-dimensional hydrogels has become increasingly common - a development that is driven by the rapidly advancing fields of tissue engineering and biofabrication. Currently, the most widely used hydrogels for 3-D cell culture are extracellular matrix (ECM)-based natural biopolymers such as collagen, fibrin, or Matrigel (a mixture of different extracellular matrix components). These biopolymers form soft, porous networks with elastic moduli on the order of 100 Pa and pore sizes on the order of microns.^1–3^ They support the adhesion of cells, and they can be locally remodeled by the cell through mechanical forces and the secretion of matrix-degrading enzymes.^4^ These properties together lead to cell behavior that is desirable for many applications, such as a high degree of cell spreading and polarization, migration, and proliferation.

Natural ECM-based biopolymer hydrogels, however, have several disadvantages. Their extreme softness and slow polymerization speed make them unsuitable for most biofabrication applications, and since the extracellular matrix components are extracted from animal sources, they can show large batch-to-batch variability, leading to poorly reproducible cell behavior. To overcome these problems, several alternative natural or synthetic hydrogels have been developed and tested, most prominently among them alginate-based, hyaluronic acid (HA)-based and polyethylene glycol (PEG)-based hydrogels.^5^ In their unmodified form, alginate, HA and PEG-based hydrogels are relatively stiff (order of 10s of kPa), their pore size is small (order of nm), they do not support strong (integrin-based) cell adhesion, and they cannot be degraded by cell-secreted proteinases.^6^ As a result, cells embedded in these hydrogels, although they can survive for weeks, remain spherical, migrate poorly, and proliferate slowly.

Cell adhesion, spreading, migration, and proliferation in PEG-, HA- and alginate-based hydrogels can be significantly improved by lowering the stiffness to below 10-20 kPa, and by adding adhesive and proteinase-sensitive components to these hydrogels.^6–9^ Lowering of the stiffness can be achieved by reducing the density of crosslinkers and by reducing the molecular weight (chain length) of the polymer components.^10^ A side-effect of these modifications is the emergence of viscoelastic behavior: When these hydrogels are deformed, the resulting mechanical stresses relax over time, and this relaxation process is faster and becomes more pronounced with decreasing molecular weight and crosslinker density.^10^

Recent reports have demonstrated that cells embedded in viscoelastic hydrogels spread and proliferate better when the hydrogels show a more pronounced stress relaxation^10^ and/or a faster stress relaxation^10,11^. The biological effects of a more pronounced stress relaxation have been explained by a greater malleability of the matrix, while the effects of a faster stress relaxation have been explained by an enhanced integrin clustering and adhesion through the formation of integrin-associated catch bonds.^12,13^

The aim of this study was to explore how desirable cell behavior such as cell spreading, migration, and proliferation depends on the time constant versus the amplitude of the viscoelastic relaxation process. For these experiments, we used alginate-based hydrogels with 1 % transglutaminase-cross-linked gelatin as an adhesive ligand. To modulate the viscoelastic relaxation behavior, we altered the concentration of cross-linking Ca^2+^ ions, and we reduced the polymer chain length and density of cross-linking sites by oxidizing alginate to alginate di-aldehyde (ADA). We also introduced a pre-crosslinking step prior to mixing-in cells and final crosslinking, which previously has been shown to lead to better cell spreading and viability^14^ and which - as we show in this study - leads to more pronounced viscoelastic relaxation.

In total, we generated 9 hydrogels with roughly similar Young’s modulus of around 1 kPa but with varying degrees of viscoelastic relaxation amplitude and relaxation speed. Our data suggest that cells respond most prominently and consistently to the viscoelastic relaxation amplitude with better spreading, faster migration, and higher proliferation. Cell behavior was also correlated with the time constant of the relaxation process, but this correlation was less pronounced and may be a spurious consequence of the fact that time constant and amplitude of stress relaxation in a material are often coupled. In conclusion, we suggest that the relaxation amplitude is a powerful design parameter for optimizing cell behavior in 3-D hydrogels.

## Experimental

### Hydrogel composition

Viscoelastic hydrogels for 3-D cell culture were based on CaCl_2_-crosslinked sodium alginate (Vivapharm PH163S2, JRS PHARMA GmbH & Co. KG, Rosenberg, Germany) with an average molecular weight of 220 kDa. For cell adhesive functionalization, alginate was either blended with gelatin (in the following referred to as Alg-GEL), or the alginate was oxidized to alginate di-aldehyde (ADA) following the approach described in ^15^, with slight adaptations. Gelatin was then covalently bound to ADA via a Schiff’s base reaction (in the following referred to as ADA-GEL).^15^ We also tested a mixture of ADA-GEL and alginate (in the following referred to as Alg-ADA-GEL). Gelatin not bound to ADA was crosslinked with 5 % (w/v) microbial transglutaminase (mTG, Activa^®^ WM, Ajinomoto Foods Europe, Germany), which was mixed and applied together with the CaCl_2_ crosslinking solution.

The oxidation process reduced the molecular weight of alginate to 120 kDa and also reduced the fraction of homopolymeric blocks of consecutive α-L-guluronate residues (GG-blocks)^16^, both of which leads to lower stiffness of the cross-linked ADA hydrogel compared to the non-oxidized alginate hydrogel for the same polymer concentrations and crosslinking solutions.^16^ Thus, to obtain hydrogels with roughly similar Young’s moduli, different concentrations of CaCl_2_ crosslinking solutions were used for ADA-GEL (100 mM), Alg-ADA-GEL (30 mM) and Alg-GEL (10 mM).

We also explored three different degrees of pre-crosslinking of the alginate or ADA copolymer prior to mixing-in cells and exposing the gels to the final crosslinking solution. Pre-crosslinking causes a heterogeneous spatial distribution of cross-linkers and - as we demonstrate in this study - influences the viscoelastic relaxation behavior of the hydrogels. Pre-crosslinking is achieved by mixing CaCO_3_ particles into the alginate or ADA solution, and releasing Ca^2+^ ions by reducing the pH with Glucono-δ-Lactone (GDL) following the approach described in ^14^.

The combination of three different hydrogels (ADA-GEL, Alg-ADA-GEL or Alg-GEL) with three degrees of pre-crosslinking (no, medium or high) resulted in nine different test conditions. Table 1 lists their compositions and the concentrations of all constituents and cross-linking solutions.

**Table 1:** Overview of final concentrations of polymer components, CaCO_3_ and Glucono-δ-lactone (GDL) pre-crosslinking solution, and CaCl_2_ and microbial transglutaminase (mTG) crosslinking solution for different hydrogel compositions.

| No. | ADA | Alg | GEL | Pre-crosslinking solution |  | Crosslinking solution |  |
| --- | --- | --- | --- | --- | --- | --- | --- |
|  | (% w/v) | (% w/v) | (% w/v) | CaCO<sub>3</sub> (mM) | GDL (mM) | CaCl<sub>2</sub> (mM) | mTG (% w/v) |
| 1 | 1.0 | 0 | 1.0 | 0 | 0 | 100 | 5 |
| 2 | 0.5 | 0.5 | 1.0 | 0 | 0 | 30 | 5 |
| 3 | 0 | 1.0 | 1.0 | 0 | 0 | 10 | 5 |
| 4 | 1.0 | 0 | 1.0 | 10 | 20 | 100 | 5 |
| 5 | 0.5 | 0.5 | 1.0 | 10 | 20 | 30 | 5 |
| 6 | 0 | 1.0 | 1.0 | 10 | 20 | 10 | 5 |
| 7 | 1.0 | 0 | 1.0 | 20 | 40 | 100 | 5 |
| 8 | 0.5 | 0.5 | 1.0 | 20 | 40 | 30 | 5 |
| 9 | 0 | 1.0 | 1.0 | 20 | 40 | 10 | 5 |

To prepare stock solutions, Alginate PH163S2 (Alg), alginate dialdehyde (ADA), or both components mixed at a mass ratio of 1:1 (Alg-ADA) were dissolved in Dulbecco’s Phosphate Buffered Saline (DPBS) at a total polymer concentration of 2.25 % (w/v). Gelatin (Type A, from porcine skin, Bloom 300, Sigma-Aldrich) was dissolved at 2.25 % (w/v) in Dulbecco’s Modified Eagle’s Medium ((DMEM) with phenol red, 4.5 g/l glucose supplemented with 10 % (v/v) bovine calf serum (BCS), 1 U/l penicillin-streptomycin (PS), 4 mM L-Glutamine, and 1 mM sodium pyruvate). Alginate stock solutions were then passed through 0.45 μm filters, and gelatin stock solutions were passed through 0.22 μm filters (Carl Roth, Karlsruhe, Germany). CaCO_3_ particles (EMSURE CAS 471-34-1, Merck, Darmstadt, Germany) were dry sterilized at 180 °C for two hours. 180 mM or 360 mM GDL solutions (CAS 90-80-2, Merck KGaA, Darmstadt, Germany) were prepared in DMEM prior to use and passed through a 0.22 μm filter.

To fabricate the hydrogel matrices, alginate-based and gelatin solutions were mixed at a volume ratio of 1:1 at 37 °C. Afterwards, pure DMEM was added to samples without pre-crosslinking to obtain the final concentrations of 1 % (w/v) gelatin and 1 % (w/v) alginate base, and mixed for 10 minutes at 37 °C with a magnetic stirrer. To prepare samples of pre-crosslinking, the gelatin/alginate-base solution was slowly added to sterile CaCO_3_ particles and mixed for another 20 min. For pre-crosslinking, 1 part of GDL solution was slowly added to 8 parts of the gelatin/alginate-base/CaCO_3_ particle mixture. Pre-crosslinked solutions were then stirred for 3 hours at 37 °C.

### Mechanical characterization of hydrogel matrices

For measuring Young’s moduli, cylindrical samples (2 mm height, 12 mm diameter) were compressed with an Instron 5967 universal testing machine (Instron, Norwood, Massachusetts, US) with a 100 N maximum force load cell (Cat# 2530-100N). Cylindrical samples were prepared as follows. First, a filter paper was soaked with the crosslinking solution and placed in a 16 cm diameter Petri dish. Next, a 2 mm thick silicone rubber sheet (ECOFLEX^®^ SERIE, KauPo Plankenhorn e.K., Germany) containing ten 12 mm diameter holes was placed on top of the filter paper. The holes were filled with 0.255 ml of freshly prepared hydrogel solution using a displacement pipette. After letting the hydrogels settle for 10 min at 4 °C, another filter paper soaked with the crosslinking solution was placed on top of the molds, and the Petri dish was flooded with the crosslinking solution. After 10 min, the crosslinking solution was removed, the silicone rubber sheet was carefully peeled off, and the hydrogel samples were transferred to Hanks’ Balanced Salt Solution (HBSS) for 10 minutes and finally to DMEM + 1 % v/v penicillin-streptomycin (PS) solution.

Samples were incubated at 37 °C in DMEM + 1 % v/v PS for 1 day, 3 days, 7 days and 10 days. Prior to measurements, the samples were equilibrated at room temperature for at least 20 min. Measurements were performed in DMEM at room temperature.

Samples were compressed with a constant speed of 0.05 mm/s up to a maximum compressive strain of 15 %. For each condition and time point, at least 3 samples were prepared and characterized.

Stress relaxation was measured with a 12 mm parallel plate rheometer (Discovery-Hybrid-Rheometer-3, TA Instruments Ltd., New Castle, Delaware, USA). Samples were prepared and incubated for 1 day as described above. For measurements, samples were placed into a glass petri dish and submersed in DMEM at 37 °C. A 12 mm parallel plate geometry was used to exert a uniaxial compressive strain at a speed of 1 mm/min up to a strain of 5 %, which was in the linear-elastic range for all tested materials. After reaching a compressive strain of 5 %, stress relaxation was recorded over a time-period of 660 s. For each condition, 4 samples were prepared and characterized. The stress relaxation was normalized to the maximum stress at the end of the loading phase, and the relaxation curves of the 4 samples of each type of hydrogel were averaged.

### Cell culture

NIH-3T3 tdTomato-farnesyl cells (stably expressing red-fluorescent tandem dimer (td) tomato protein carrying a CAAX-sequence that upon farnesylation recruits the protein to the cell membrane) were generated by lentiviral transduction of the tdTomato-farnesyl-5 reporter construct as described in ^17^. NIH-3T3 tdTomato-farnesyl cells (hereinafter referred to as NIH-3T3 tdTomato cells) were in 75 cm² flasks with phenol-red containing Dulbecco’s Modified Eagle’s Medium (DMEM), supplemented with 4.5 g/l glucose, 10 % (v/v) bovine calf serum (BCS), 1 U/l Penicillin-Streptomycin (PS), 4 mM L-Glutamine and 1 mM sodium pyruvate. Cell culture flasks were incubated at 37 °C, 5 % CO_2_ and 95 % humidity, and cells were passaged every second to third day using 0.25 % Trypsin-EDTA (Thermo Fisher Scientific, Germany) at a ratio between 1:6 and 1:8, depending on the initial cell density.

### Hydrogel embedding

Experiments for 3-D cell culture were conducted in 24-well cell culture plates (Sarstedt, Germany). Each well was first filled with 200 μl of Alg-GEL 1.0%/1.0% solution (solution #3) without cells, and gelled at 4 °C for at least 20 min. This layer was applied to avoid direct attachment of the cells to the bottom of the well plate. In the meantime, the cells were detached from cell culture flasks with 0.25 % Trypsin-EDTA (Thermo Fisher Scientific, Germany) and counted with a Lobauer counting chamber (Neubauer-improved, Paul Marienfeld, Lauda-Königshofen, Germany). Cells were centrifuged at 300 rcf for 5 minutes, and the cell pellet was mixed with the polymer solution using a positive displacement pipette to a final concentration of 40,000 cells/ml for cell migration experiments, or 100,000 cells/ml for cell proliferation experiments. 200 μl of the cell-polymer solution was added to each well. The cross-linking solution was then applied with a spray bottle until the sample was covered with at least 1 ml of cross-linking solution. After 10 min, the crosslinking solution was removed, and samples were washed with Hanks’ Balanced Salt Solution (HBSS) before adding the complete cell culture medium. Cells were incubated at 37 °C, 5 % CO_2_ and 95 % humidity for up to ten days. Cell culture medium was replaced three times per week.

### Colorimetric cell proliferation assay (WST-8)

Cell proliferation was estimated based on a colorimetric NADH activity assay using the Cell Counting Kit - 8 (Sigma Aldrich – Product #96992) according to the manufacturer’s instructions with slight adaptations. For the staining solution, the WST-8 dye indicator solution was mixed with phenol-red free complete cell culture medium at a concentration of 50 μl/ml. At day 1, day 3, day 7 and day 10 of culture, cell culture medium was removed from the hydrogel samples and replaced with 300 μl staining solution per well. Samples were incubated for 4 h at 37 °C, 95 % humidity and 5 % CO_2_. Two probes of 100 μl of the supernatant from each well were transferred into two separate 96-wells of a plate for technical repeats. Absorbance was measured at a wavelength of 450 nm using a plate reader (FLUOstar Omega, BMG Labtech, Ortenburg, Germany). Supernatants from cell-free hydrogel samples that underwent the same experimental procedure were used as control. The absorbance of the control sample was subtracted from the matching cell-containing sample. For each condition, 5 hydrogel samples were measured.

### Cell migration

Cell migration was measured at day 1, day 3, day 7 and day 10 using time-lapse imaging. Before imaging, samples were stained with 2.5 μg/ml Hoechst 33342 (Sigma-Aldrich, #14533) for 30 min at room temperature, washed and covered with 1 ml complete cell culture medium. 31 bright field z-stack images over a depth of 300 μm and a z-distance of 10 μm were taken every 10 min for a total duration of 12 h using a motorized microscope (ASI, Eugene, OR, USA) equipped with a 10x 0.3 NA objective (Olympus) and a CMOS camera (Lumenera Infinity 3-6URM, Teledyne Lumenera, ON, USA) inside a cell culture incubator (37 °C, 5 % CO_2_ and 95 % humidity). Additionally, fluorescent (391/450 nm) z-stacks were imaged (every 10 min over 24 h). Subsequently, automatic tracking of the x/y-coordinates of the cells’ fluorescent nuclei in an image series of maximum intensity z-stack projections was performed, using a Clickpoints-based Python script (https://github.com/fabrylab/tracking_andreas). At least three and up to six fields of view were imaged at each time point. Cell trajectories shorter than 4 h were excluded from subsequent analysis. Table 2 shows the total amount of evaluated cell trajectories per condition.

**Table 2:** Overview of total number of evaluated cell trajectories per condition

| Material | Pre-crosslinking degree | Day 1 | Day 3 | Day 7 | Day 10 |
| --- | --- | --- | --- | --- | --- |
| ADA-GEL | no | 87 | 149 | 128 | 298 |
|  | medium | 71 | 159 | 337 | 327 |
|  | high | 117 | 394 | 203 | / |
| Alg-ADA-GEL | no | 76 | 62 | 99 | 68 |
|  | medium | 75 | 119 | 110 | 89 |
|  | high | 51 | 205 | 243 | 358 |
| Alg-GEL | no | 61 | 47 | 37 | 108 |
|  | medium | 51 | 65 | 101 | 97 |
|  | high | 41 | 37 | 54 | 92 |

Cells were classified as motile if they migrated at least 20 μm away from their starting point (Euclidean distance) during the total observation period. The mean cell migration speed was calculated from the Euclidean distance of cell coordinates between two images taken 60 minutes apart, averaged over the entire track length.

### Cell morphology

Bright-field and fluorescent images (554/581 nm) for the evaluation of cell morphology were taken prior to starting the cell migration time-lapse recording, at the same positions, except that each stack contained 60 single images with a z-distance of 5 μm.

Depending on the turbidity of the samples, two different approaches were used. In the case of transparent samples (non- and medium-pre-crosslinked Alg-ADA-GEL and Alg-GEL samples), maximum-intensity projections of the fluorescence image stacks were contrast-enhanced, bandpass-filtered to remove the background, and binarized with Fiji ImageJ using a manually adjusted image-specific threshold. Subsequently, binary images showing cell shapes were analyzed using the Fiji ImageJ software plugin “analyze particles”. In the case of samples that showed high turbidity either due to pre-crosslinking or cell crowding) (all ADA-GEL samples and high-pre-crosslinked Alg-ADA-GEL and Alg-GEL), no maximum-intensity projection was performed, and cells were manually segmented from the bright field images using the ClickPoints software plugin “cell measurement”.^18^ The cell circularity C was computed as

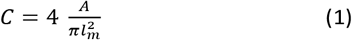

with *A* being the cell area and *l*_*m*_ the major axis length (endpoints of the longest line that can be drawn through the segmented cell shape). Circularity is unity for circular cells and tends towards zero for highly elongated cells. This definition of circularity was chosen because it is insensitive to cell shape irregularities but sensitive to the presence of cell protrusions. The circularity was calculated for at least 30 individual cells per condition. At least two and up to seven fields of view were evaluated for each sample, depending on the number of cells per image. Three individual samples were measured for each condition and time point.

### Statistical analysis

Rank correlations were computed to test for correlations between biological (motility, migration speed, circularity and proliferation) and mechanical (relaxation amplitude, relaxation time and Young’s modulus) data. For a guide-to-the-eye visualization of the relationship between mechanical and biological responses in the case of significant correlations (p < 0.05), a logistic function in the form of

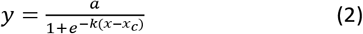

with amplitude *a*, midpoint *x*_*c*_ and steepness *k*, was fitted to the data.

## Results and discussion

### Young’s moduli

Non-pre-crosslinked hydrogels of different composition (ADA-GEL, Alg-ADA-GEL and Alg-GEL) that were crosslinked with different concentrations of CaCl_2_ had similar Young’s moduli in the range of 1.4-1.9 kPa after 1d of incubation (*Figure 1 a*). This confirms that lowering the CaCl_2_ concentrations of the crosslinking solution was effective to counterbalance the influence of an increasing molecular weight of the polymer phase in the different hydrogels.

**Figure 1:**
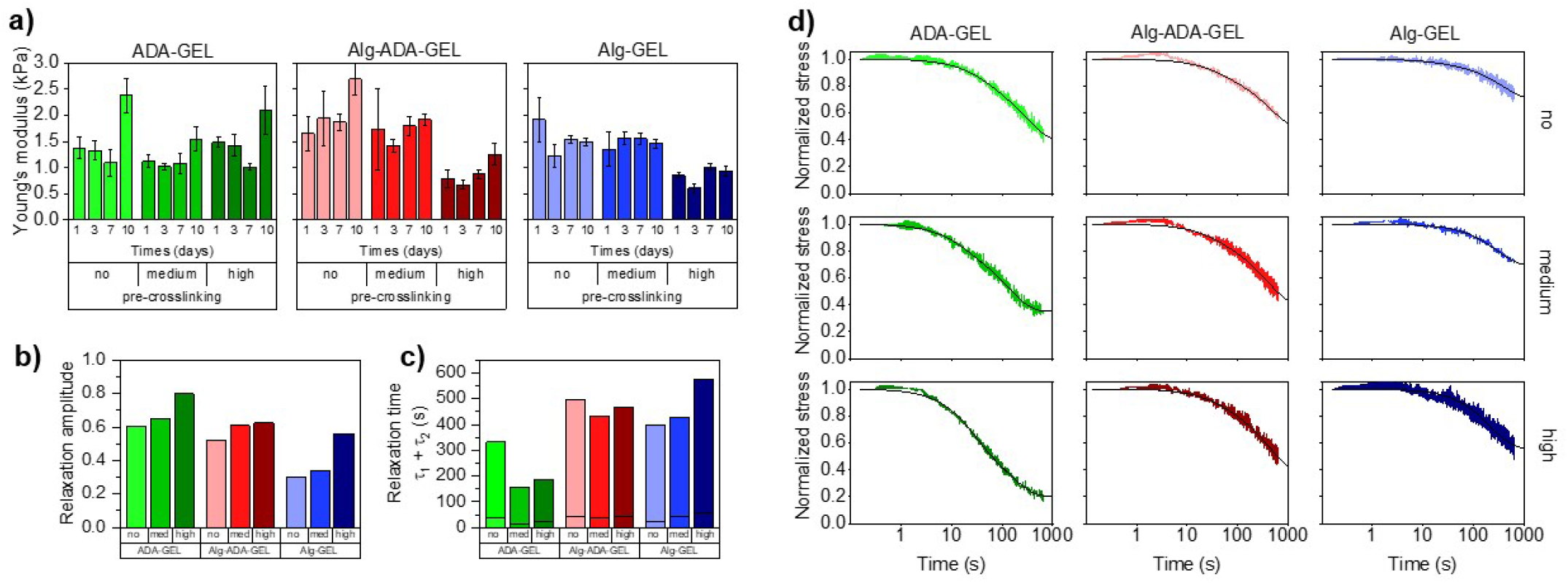
a) Young’s moduli (mean ± SD, n≥3) measured after 1, 3, 7 and 10 days of incubation for a compressive strain of 5 %. b) Stress relaxation amplitude (*σ*1 + *σ*2) and c) characteristic time constants (*τ*1 (bottom) and *τ*2 (top), separated by horizontal lines). d) Stress-relaxation behaviour of non-, medium- and high-pre-crosslinked ADA-GEL, Alg-ADA-GEL and Alg-GEL samples after 1 day of incubation. Hydrogels were compressed to a compressive strain of 5 %, which was then held constant for 600 s. Stress was normalized to the maximum value when the final compressive strain of 5 % was reached and averaged (n = 4). Lines represent the fit of Eq. 3 to the data.

After longer incubation times, the Young’s modulus of the hydrogels changed slightly, but did neither consistently increase or decrease. For ADA-GEL and Alg-ADA-GEL samples but not for Alg-GEL samples, we found an increase in the Young’s modulus after 10 days of incubation in cell culture medium. Currently we do not know the origin of this increase. Microscopic inspection did not reveal any structural changes, and - as we present below - cell behavior after a culture time of 10 days did also not show any noticeable qualitative alterations compared to 7 days of culture.

A medium degree of pre-crosslinking had no noticeable effect on the Young’s modulus. A high degree of crosslinking lowered the Young’s modulus of the Alg-ADA-GEL and Alg-GEL hydrogels by approximately 50 %, but had no effect in ADA-GEL hydrogels. After longer incubation times of up to 10 days, the Young’s modulus of the pre-crosslinked hydrogels also changed slightly but not consistently in either directions, similar to the behavior of the samples that were not pre-crosslinked. This is consistent with the notion that the hydrogels did not substantially degrade or change structurally over time.

### Stress relaxation behavior

The stress relaxation *σ*(*t*) over time *t* normalized to the stress *σ*_0_ at the beginning (*t* = 0) of a rapid strain increase by 5 % was well described by the superposition of two exponential functions

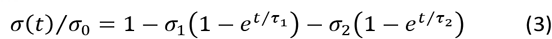

with *σ* being the amplitude and *τ* being the time constant of the individual exponentials. The total stress relaxation amplitude can be expressed as *σ*_1_ + *σ*_2_.

When comparing the relaxation behavior of the nine hydrogels, we observed the following trends: Hydrogels with increasing degree of pre-crosslinking and with increasing amount of oxidized alginate dialdehyde (ADA) displayed a larger relaxation amplitude (*Figure 1 b*). The relaxation-promoting effect of pre-crosslinking was particularly pronounced in the ADA-GEL hydrogels. The speed of the fast relaxation process (characterized by the time constant *τ*_1_) was around 15-60 s, and the speed of the slow relaxation process (characterized by the time constant *τ*_2_) was between 150 s and 600 s. Both tended to be faster for gels that displayed a larger relaxation amplitude. The correlation between the relaxation amplitude *σ*_1_ + *σ*_2_ and the relaxation time constants, however, was not statistically significant (p > 0.05).

### Cell proliferation

Cell proliferation of NIH-3T3 tdTomato cells embedded in the different hydrogel samples was measured after 1, 3, 7 or 10 days of cell culture using a colorimetric NADH activity assay. This assay measures the absorbance of the dye solution (decadic logarithm of the ratio of transmitted to incident light intensity) at a wavelength of 450 nm, which scales with cell number.^19^ We subtract from this value the absorbance of a dye solution taken from cell-free hydrogel samples. For measuring the proliferation of cells cultured in a 3-D hydrogel, we adapted the manufacturer’s protocol, which is optimized for 2-D culture conditions, and used a larger dye volume (300 μl dye solution for a 400 μl hydrogel sample) and longer exposure time (4 h). We consider absorbance values below 0.05 to be below the noise floor and values above 0.8 to be outside the dynamic range of the method.

Proliferation was marginal in Alg-GEL hydrogels and remained below the noise floor even after a culture time of 10 days (Figure 2 a). In Alg-ADA-Gel hydrogels, we observed a small increase in cell number over time, which became more pronounced in the pre-crosslinked samples. By contrast, in the ADA-Gel hydrogels, we observed a strong increase in cell density over time, which also became more pronounced in the pre-crosslinked samples. After 10 days of culture, individual cells in the highly pre-crosslinked ADA-Gel hydrogels could no longer be reliably discerned due to overcrowding (*Figure 2 b*). This overcrowding was not detected by the colorimetric NADH activity assay as the corresponding increase in absorbance was outside the dynamic range.

**Figure 2:**
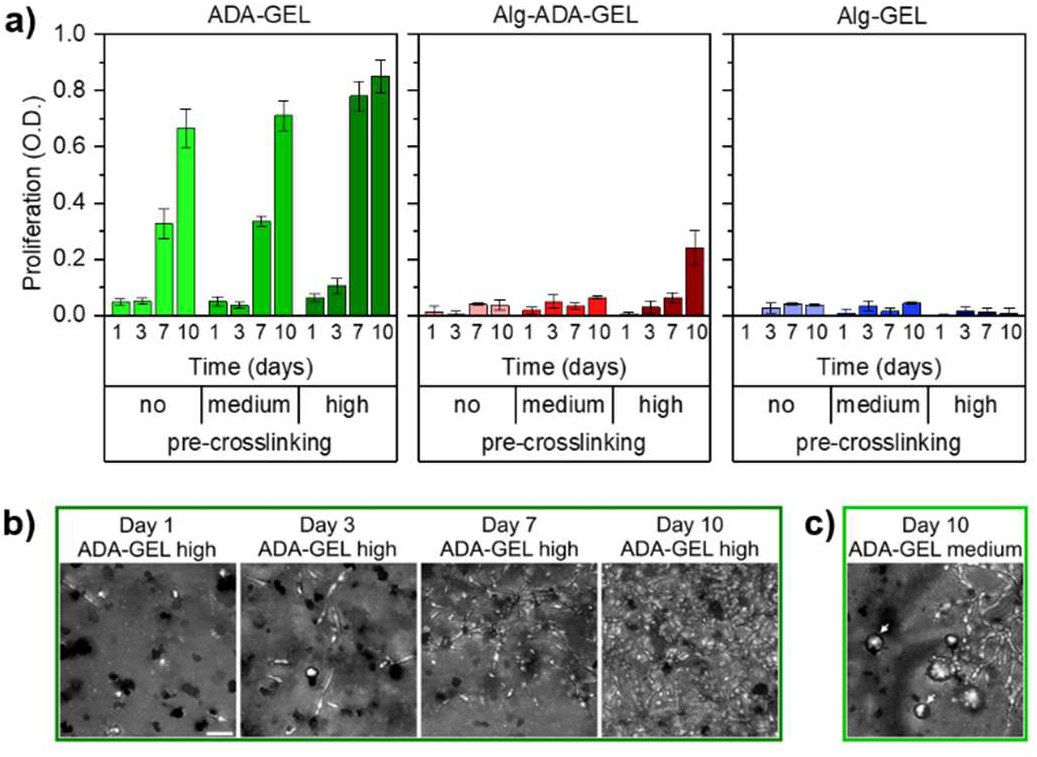
a) Colorimetric cell proliferation assay (WST-8) of NIH-3T3 tdTomato cells. Cells were embedded in non-, medium- and high-pre-crosslinked ADA-GEL, Alg-ADA-GEL and Alg-GEL samples for 1, 3, 7 and 10 days. Data were obtained from n=5 experiments, each with n=2 technical repeats. All data are displayed as mean ± SD. b) Representative maximum intensity projection images to visualize the increase in cell numbers within high-pre-crosslinked ADA-GEL (day 1,3,7,10). Representative images for all conditions can be found in Fig. ESI2. c) Representative maximum intensity projection image of a medium-pre-crosslinked ADA-GEL after 10 days shows inhomogeneous behavior characterized by the presence of elongated cells together with spherical cell agglomerates (white arrows). Scale bar = 50 μm.

### Cell migration

Cell migration of single cells was quantified from the trajectories of 2-D projected (x,y) cell positions recorded every 10 min over a total observation time of 24 h. In particular, we calculated the 1 h migration speed as the Euclidean distance between all (x,y) positions of the same cell that were 60 min apart, averaged over 24 h (Figure 4).

**Figure 3:**
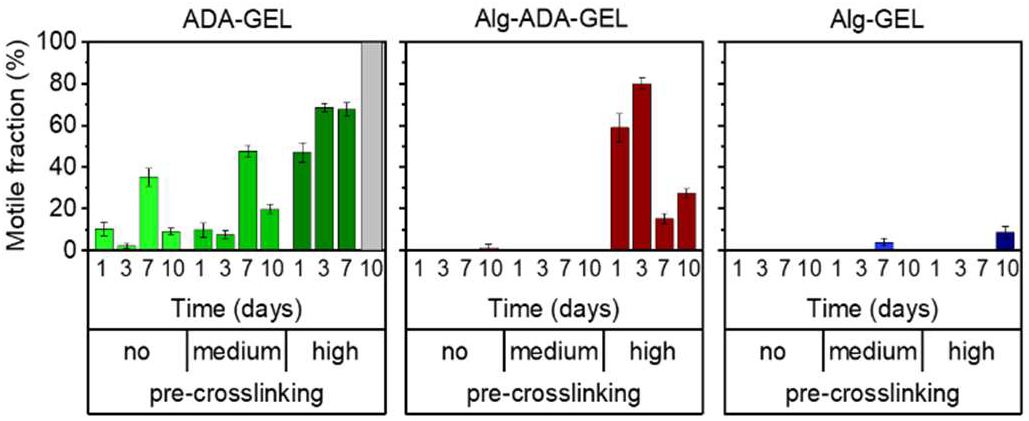
Fractions of motile cells within non-, medium- and high pre-crosslinked ADA-GEL, Alg-ADA-GEL and Alg-GEL. Cells were classified as motile when they migrated by more than 20 μm in 24 h. Graph shows mean ± se from 3-6 FOV of 3 samples (n≥37 analyzed cells).

**Figure 4:**
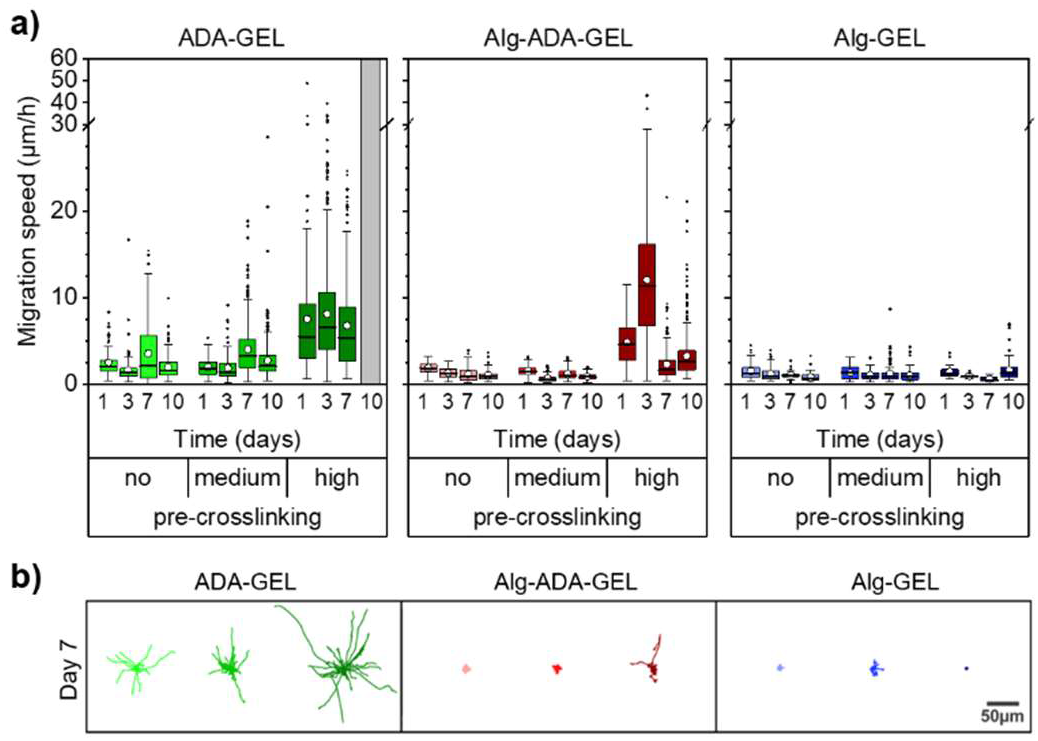
a) Migration speed of tracked single cells embedded in non-, medium- and high-pre-crosslinked ADA-GEL, Alg-ADA-GEL or Alg-GEL hydrogels. White circles represent mean, horizontal lines the median, whiskers indicate the 1.5 interquartile range, and outliers are shown as black points from 3-6 FOV of 3 samples (n≥37 analyzed cells). Cells could not be tracked at day 10 in high-pre-crosslinking ADA-GEL due to excessive crowding. b) Rose-plots of 36 exemplary cell tracks (4 h duration) for each condition at day 7.

Cells did not or only marginally migrate in Alg-GEL hydrogels, regardless of pre-crosslinking and culture time. Similarly, in non-pre-crosslinked and medium pre-crosslinked Alg-ADA-GEL hydrogels, cells did not migrate appreciably. In high pre-crosslinked Alg-ADA-GEL hydrogels, however, cells started to migrate already after 1 day of culture. The migration speed was largest after 3 days in culture and reached values of more than 10 μm/h on average. For longer culture times, migration speed decreased to below 5 μm/h on average, but a substantial fraction of the cells migrated with speeds > 10 μm/h.

In all ADA-GEL hydrogels, cells migrated well, with an average migration speed that remained approximately constant over 10 days. With higher pre-crosslinking concentration, migration speed increased and reached values above 5 μm/h, with a substantial fraction of the cells migrating faster than 20 μm/h.

### Motile fraction

While some cells migrated large distances during the observation period, other cells did not migrate at all. To analyze this behavior, we grouped the cells into a motile and immotile fraction, depending on whether a given cell migrated by more than 20 μm over a total observation period of 24 h.

The behavior of the motile fraction corresponded roughly to the behavior seen for the migration speed (Figure 3): The motile fraction was negligible for all Alg-GEL and for non- and medium-pre-crosslinked Alg-ADA-GEL hydrogels. In the high-pre-crosslinked Alg-ADA-GEL hydrogels, the motile fraction was largest during the first 3 days of culture (>50 %) and then declined to about 20-30 %. In the ADA-GEL hydrogels, the motile fraction increased with pre-crosslinking concentration and reached values above 60 % for highly pre-crosslinked hydrogels.

### Cell morphology

Non-motile cells typically had a circular shape, whereas motile cells were mostly elongated. We therefore quantified the circularity of the x,y-projected cell shapes in the different hydrogels. Circularity values near unity indicate a nearly circular shape, whereas values near zero indicate elongated cell shapes. The circularity of the cells in the different hydrogels roughly followed the trend seen for the migration speed and the motile fraction, however with some noticeable differences (Figure 5). For example, cells in the highly pre-crosslinked Alg-Gel hydrogels started to become elongated after 10 days of culture, even though the motile fraction and the migration speed was still small. Furthermore, despite the decrease in the migration speed and motile fraction over time in the highly crosslinked Alg-ADA-GEL hydrogels (after 7 and 10 days of culture), the cells remained highly elongated. In the ADA-GEL hydrogels, cells monotonically elongated over time, whereas the trend for the motile fraction and migration speed was less consistent.

**Figure 5:**
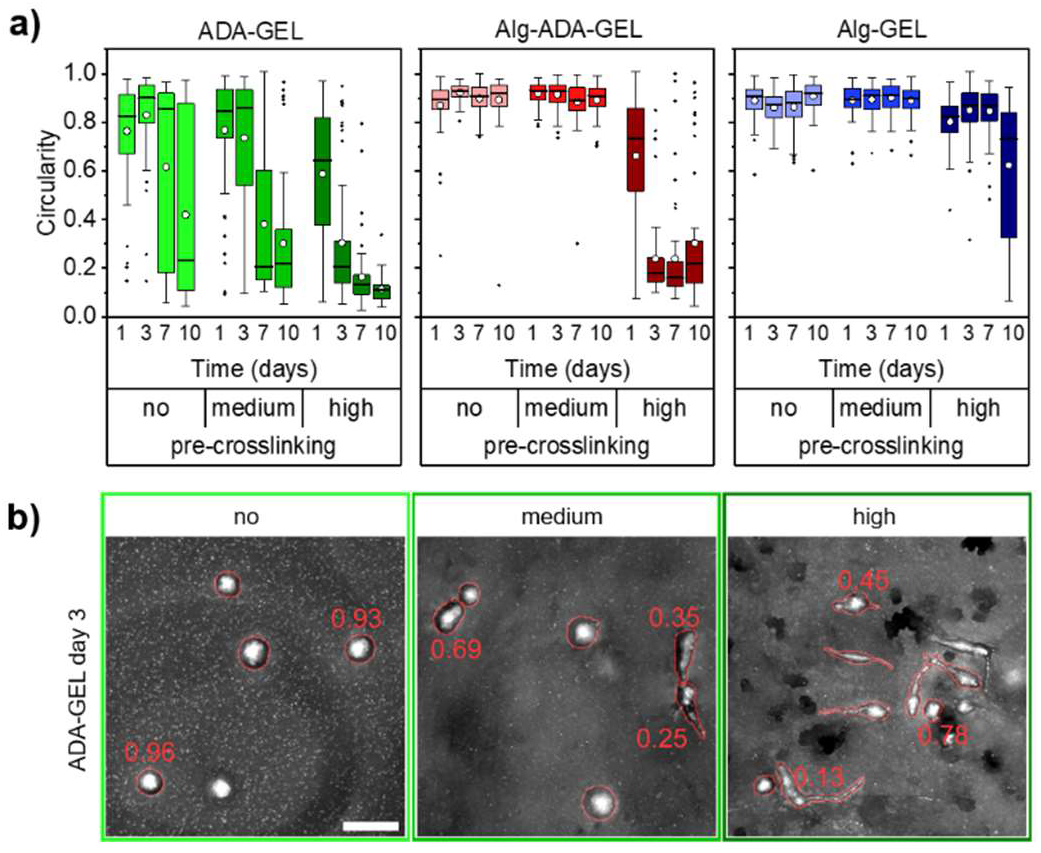
a) Circularity of NIH/3T3 tdTomato cells embedded in non-, medium- and high-pre-crosslinked ADA-GEL, Alg-ADA-GEL and Alg-GEL hydrogel samples for 1, 3, 7 and 10 days. White circles represent mean, horizontal lines the median, whiskers indicate the 1.5 interquartile range, and outliers are shown as black points from at least 2 FOV of 3 samples (n≥30 analyzed cells). b) Representative maximum intensity projection images of cells embedded in non-, medium- and high-pre-crosslinked ADA-GEL samples after 3 days of incubation. Numbers depict circularity values of exemplary cells. Scale bar = 50 μm.

### Correlation of stress relaxation behavior and cell behavior

Taking all cell behavior results together, cells elongated, proliferated, and migrated better in the more highly pre-crosslinked hydrogels, in particular in the ADA-Gel hydrogels. More pre-crosslinking was also accompanied by a higher amplitude of stress relaxation, and hence, we found strong relationships between relaxation amplitude versus proliferation, migration speed, motile fraction, and circularity (Figure 6). To test if these relationships were monotonic and statistically significant, we computed the Spearman rank correlation between the cell biological values (measured after 7 days and 10 days of culture) and the relaxation amplitude. We found in each case a high and statistically significant (p < 0.05) correlation of the cell biological parameters with the hydrogels’ relaxation amplitude.

**Figure 6:**
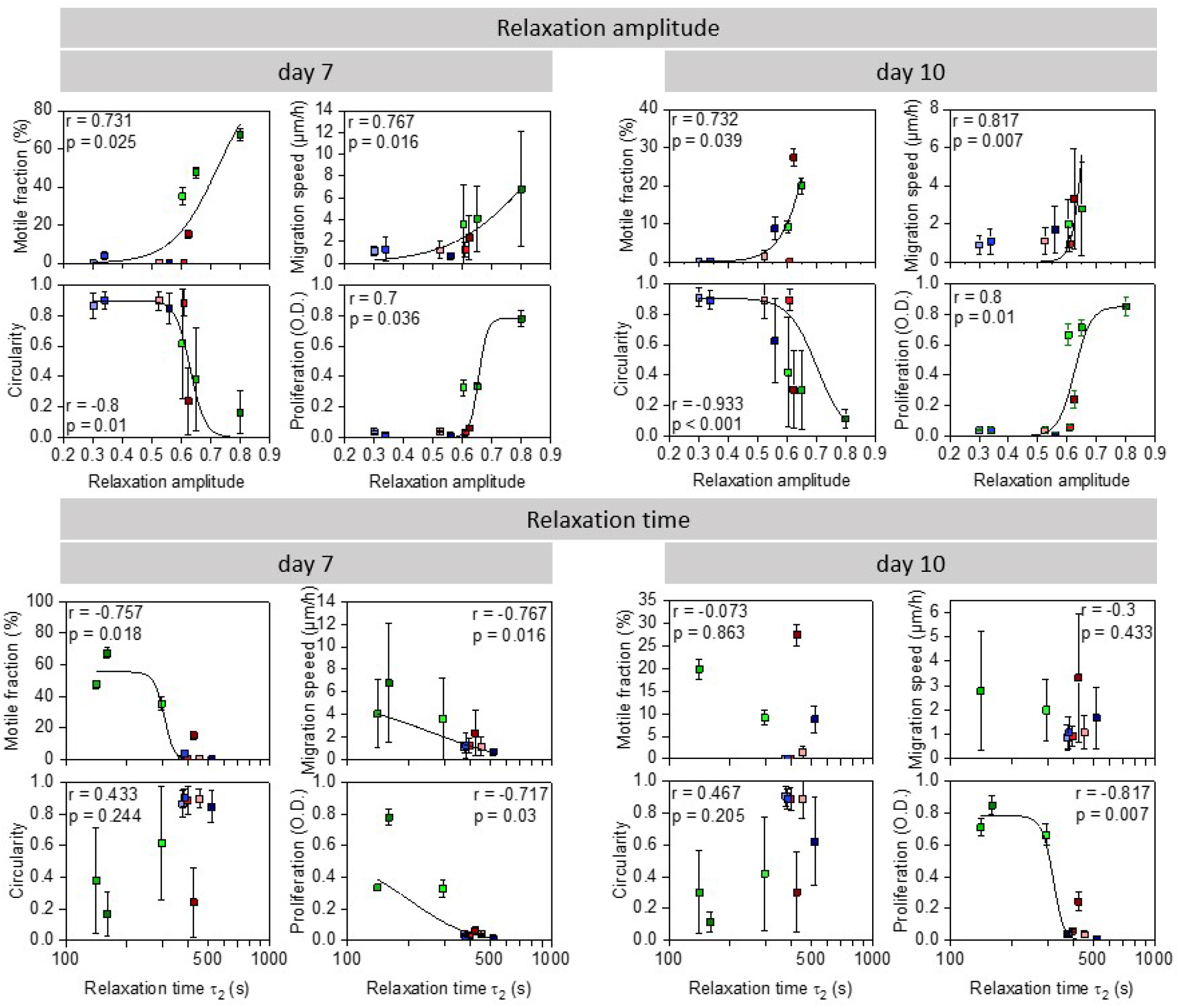
Correlation between relaxation amplitude (above) or slow time constant (*τ*_2_) (below) and motile fraction of cells, migration speed, circularity, and cell proliferation on day 7 (left) and day 10 (right). Rank correlation r and its significance value p are indicated in the graphs. Black lines are a fit of Eq. 2 to the data with statistically significant correlations as a “guide to the eye”.

We also performed the same correlation analysis of the cell biological behavior with the slow time constant (*τ*_2_) of the relaxation behavior. Generally, the correlation was less pronounced but also reached statistical significance (p < 0.05) in the case of cell proliferation (7 and 10 days), migration speed at 7 days, and the motile fraction at 7 days.

We also tested if the cell biological behavior was correlated with the Young’s moduli of the hydrogels, but we found no significant correlation (ESI1).

## Discussion

In this study, we explored the shape, migration, and proliferation of cells embedded in alginate-based hydrogels with different viscoelastic relaxation properties. We achieved a variation in the viscoelastic relaxation behavior by altering the polymer chain length (through different fractions of oxidized alginate (ADA), which has a lower molecular weight compared to alginate), and by altering the degree of polymer chain crosslinking. Different degrees of polymer chain crosslinking, in turn, were achieved by different Ca^2+^ ion concentrations of the crosslinking solution. Specifically, we increased the Ca^2+^ ion concentrations of the crosslinking solution with increasing fractions of ADA. This ensured that the Young’s moduli of the resulting hydrogels, despite their different composition, remained similar. The higher Ca^2+^ ion concentrations also ensured that the ADA-containing hydrogels remained stable and did not appreciably degrade over a time period of 10 days. Importantly, we also introduced a pre-crosslinking step prior to mixing-in the cells and prior to the final crosslinking. This pre-crosslinking has been previously shown to improve cell viability.^14^ Thus, a motivation of the present study was to test the hypothesis that the improved cell viability in pre-crosslinked alginate-based hydrogels was due to an altered viscoelastic relaxation behavior.

We found that all hydrogels showed pronounced viscoelastic stress relaxation when exposed to a rapid strain increase by 5 %. The time course of the stress relaxation was well described by a bi-exponential function (superposition of a fast and a slow exponential function), which allowed us to quantify the relaxation behavior in terms of a relaxation amplitude (the combined amplitude of the fast and the slow exponential function) and a time constant (of the slow exponential function). We found that hydrogels tended to display an increased relaxation amplitude with increasing ADA fraction and increasing degree of pre-crosslinking. The time constant of the stress relaxation also tended to increase with higher ADA fraction but did not show a clear trend with pre-crosslinking. The main finding of our study is that cells tended to elongate, migrate and proliferate better in hydrogels with higher stress relaxation amplitude.

Each cell biological parameter was significantly (p < 0.05) correlated with the stress relaxation amplitude. By contrast, the correlation of cell biological parameters with the relaxation time constants of the hydrogels was less pronounced and less consistent. Circularity, fraction of motile cells after 10 days of culture, and migration distance after 10 days of culture were not significantly correlated with the relaxation time constant, and only proliferation, fraction of motile cells after 7 days of culture, and migration distance after 7 days of culture were significantly correlated.

Previous reports suggested that cell spreading and proliferation in 3-D hydrogels increases with faster viscoelastic stress relaxation and/or with a higher amplitude of stress relaxation.^10–12,20^ While our data are in agreement with these reports, they suggest that not so much the time constant but more so the amplitude of stress relaxation is associated with higher cell spreading and cell proliferation.

To proliferate, mesenchymal cells such as the NIH-3T3 fibroblasts, used in our study, need to adhere to an extracellular matrix. Mesenchymal cells that cannot adhere tend to be circular in shape, and they proliferate poorly even if they have enough space to grow.^21^ Thus, a round morphology in mesenchymal cells typically indicates that the cells do not proliferate. Moreover, cell elongation in mesenchymal cells is associated with better and faster migration in confined 3-D environments.^22^

To spread, migrate and proliferate, cells need to displace parts of the extracellular matrix, and to do so, the matrix needs to be soft and/or degradable. The energy and pressure required in this process must be below a critical threshold or else cell proliferation stalls^23^, and cell migration comes to a halt^1,24^. Degradation can either be achieved chemically through proteolysis, or mechanically through the application of forces that are large enough to break bonds between matrix components. Since alginate-based hydrogels are not degradable through cell-secreted proteases, mechanical degradation through force application by the cell is the only option. The amplitude of stress relaxation of a material corresponds to the ability of external forces to break material bonds. Hence, it follows that the ability of cells to elongate, to migrate and to proliferate increases in hydrogels of the same initial stiffness that display a larger stress relaxation amplitude. Our data agree with this notion.

There are also convincing arguments why spreading, migration and hence proliferation also depend on the speed of stress relaxation. First, if the same degree of local bond-breakage and pore-opening at the cellular level can be achieved with the same force but in less time, the cell needs to invest less energy. Second, spreading and migration in confined space requires the cell to break symmetry and to polarize, but if the surrounding matrix is too slow to accommodate and deform accordingly, the cell will more rapidly change polarity in a different direction and - through this lack of persistence - remain round and immobile.^25^ Third, studies of cell behavior on 2-D viscoelastic substrates have reported increased migration speed on substrates with faster relaxation times.^13^ However, cells were also more circular on faster-relaxing substrates and formed only weak, nascent adhesions that do not support strong tractions that are needed for confined 3-D migration.^22,26^ Hence, the effect of stress relaxation speed on cell migration in a 3-D environment may depend on the material’s steric hindrance, which in turn depends on the elasticity, porosity, and - as argued above - on the amplitude of stress relaxation.

Although our data suggest that cell spreading, migration, and proliferation are more closely associated with the stress relaxation amplitude as opposed to the time constant of stress relaxation, we cannot establish a causal relationship. A main limitation of our study is that we cannot independently tune the amplitude versus the time constant of stress relaxation in different materials. Nonetheless, our finding that all measured cell biological parameters were significantly correlated with the stress relaxation amplitude, whereas only some cell biological parameters were correlated with the relaxation time constant, point to the relaxation amplitude as the dominant material property. In conclusion, we suggest that the stress relaxation amplitude of a hydrogel is a highly effective design parameter to tune and optimize cell behavior in 3-D hydrogels.

## Supporting information

Electronic supplementary information

Time lapse video ADA-GEL hydrogel

Time lapse video Alg-ADA-GEL hydrogel

Time lapse video Alg-GEL hydrogel

## Author Contributions

Conceptualization (all authors); Investigation (Jonas Hazur, Nadine Endrizzi); Formal Analysis (Jonas Hazur, Nadine Endrizzi, Ben Fabry); Methodology (all authors); Supervision and funding acquisition (Aldo R. Boccaccini, Ben Fabry, Dirk W. Schubert); Writing (all authors)

## Acknowledgements

We thank Ingo Thievessen and Lena Fischer for providing NIH-3T3 tdTomato cells.

Funded by the Deutsche Forschungsgemeinschaft (DFG, German Research Foundation) – Projektnummer 326998133 – TRR 225 (subprojects A01, A07, C02).

