## Supplementary material for "Stress relaxation amplitude of hydrogels determines migration, proliferation, and morphology of cells in 3-D": Electronic supplementary information

### Electronic supplementary information for the manuscript

#### Correlation of Young's modulus and cell behavior

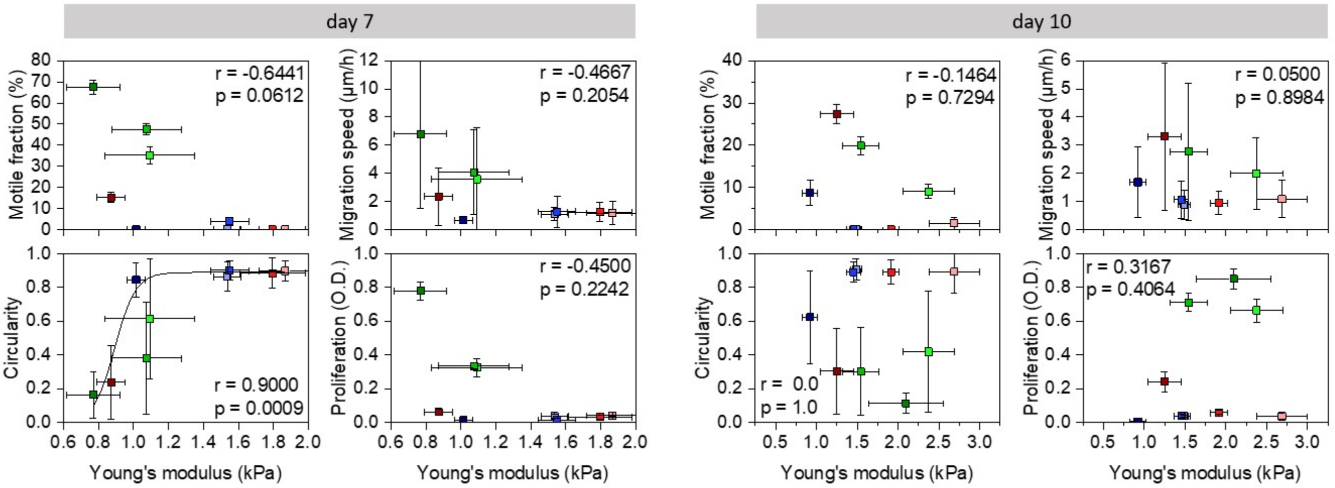


*Figure ESI 1: Correlation between Young's moduli and motile fraction of cells, migration speed, circularity, and cell proliferation on day 7 (left) and day 10 (right). Rank correlation r and its significance value p are indicated in the graphs. Black lines are a fit of Eq. 2 to the data as a "guide to the eye" (shown only for statistically significant correlations).*

#### Representative cell images and videos

Time laps videos (12.5 h duration) of maximum intensity projections of NIH/3T3 tdTomato cells embedded in non-, medium- and high-pre-crosslinked ADA-GEL, Alg-ADA-GEL and Alg-GEL hydrogel samples after 1, 3, 7 and 10 days of culture. Videos can be downloaded via the following hyperlinks for ADA-GEL, Alg-ADA-GEL and Alg-GEL. Scale bar = 50 µm.

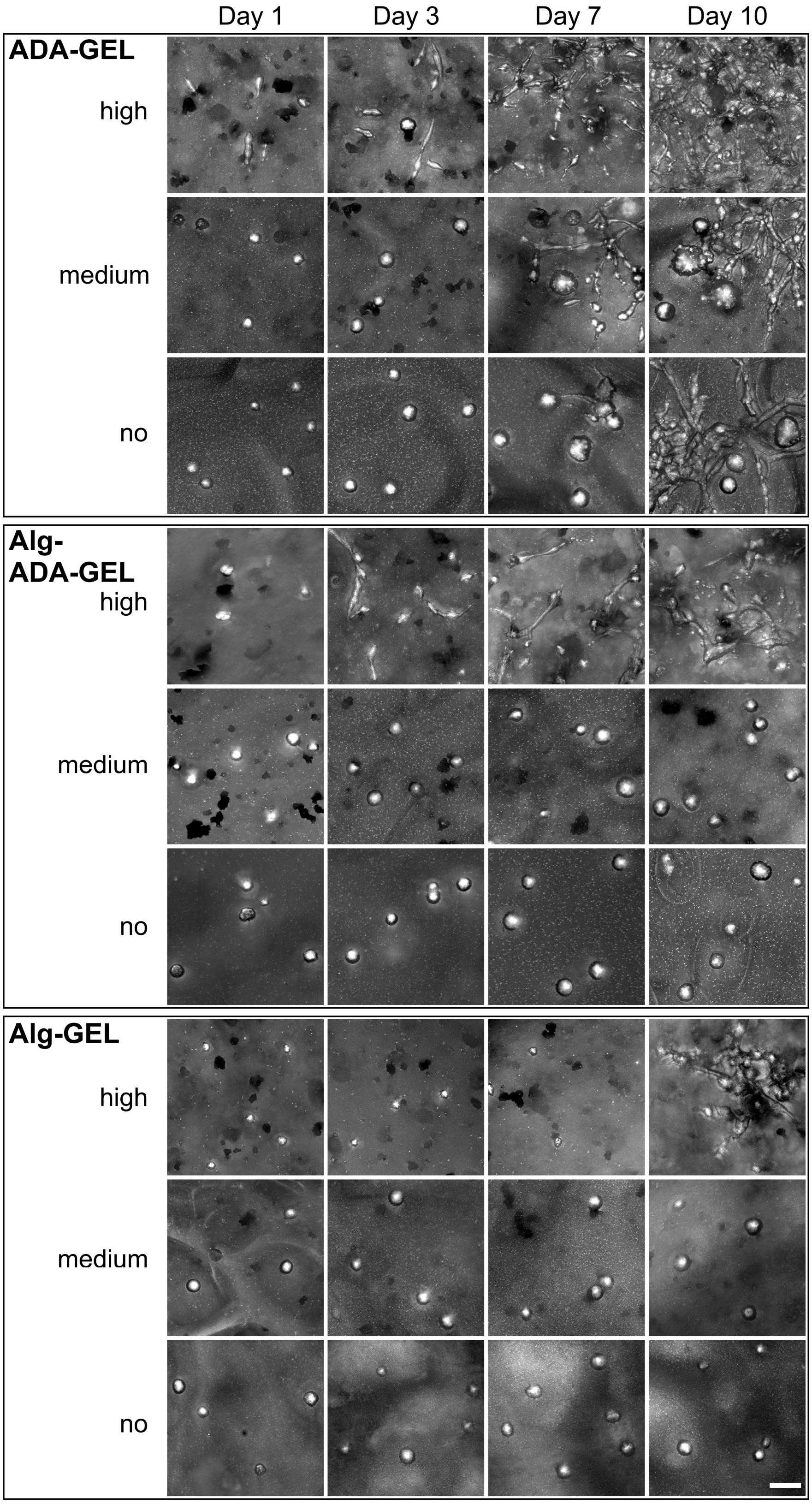


*Figure ESI 2: Maximum intensity projection images of NIH/3T3 tdTomato cells embedded in non-, medium- and high-pre-crosslinked ADA-GEL, Alg-ADA-GEL and Alg-GEL hydrogel samples after 1, 3, 7 and 10 to visualize proliferation. Scale bar = 50 µm.*

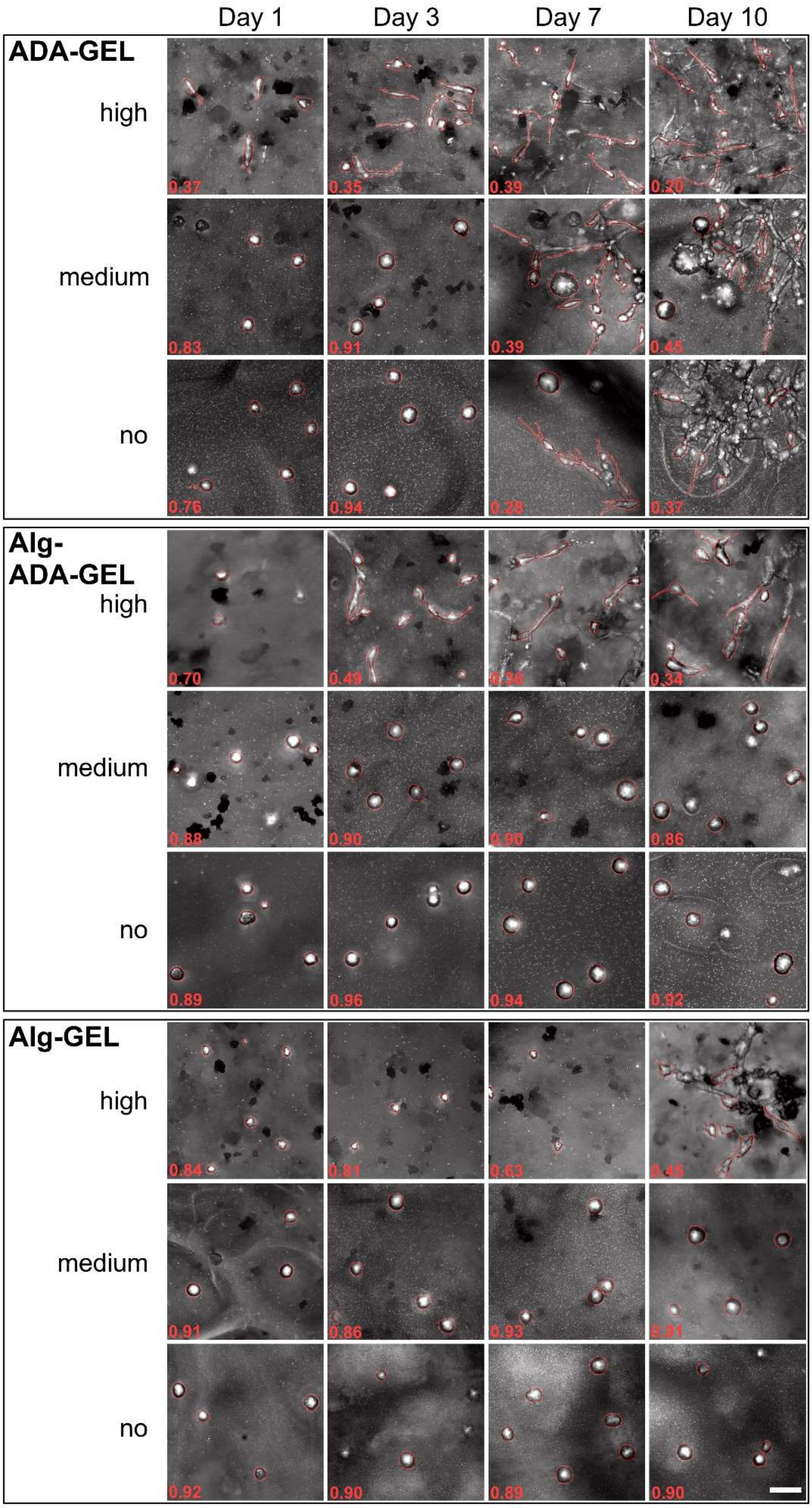


*Figure ESI 3: Maximum projection images of NIH/3T3 tdTomato cells embedded in non-, medium- and high-pre-crosslinked ADA-GEL, Alg-ADA-GEL and Alg-GEL hydrogel samples are shown for day 1, 3, 7 and 10 to visualize cell morphology. Red numbers depict mean roundness values of exemplary cells per image. Scale bar = 50 µm.*
